# Functional Suppression of Premotor Activity in a Transient Model of Motor Cortex Injury

**DOI:** 10.1101/2020.06.12.148403

**Authors:** Kevin C. Elliott, Jordan A. Borrell, Scott Barbay, Randolph J. Nudo

## Abstract

Cortical injuries (e.g. – strokes or traumatic brain injuries) can create a host of secondary events that further impair the brain’s sensory, motor, or cognitive capabilities. Here, we attempted to isolate the acute effects of the primary injury – the loss of cortical activity – on rodent motor cortex (caudal forelimb area, CFA) without the secondary effects that arise from damage to cortical tissue. We then observed the effects of this loss of activity on the rodent premotor cortex (rostral forelimb area, RFA). In anesthetized rats, CFA was temporarily inactivated with the GABA-A agonist muscimol, disrupting motor network function while leaving neural connectivity intact. Using intracortical microstimulation (ICMS) techniques, we found that CFA inactivation completely abolished ICMS-evoked forelimb movement from RFA yet spared some CFA evoked-movement. Neural recordings confirmed that neural suppression by muscimol was isolated to CFA and did not spread into RFA. We next observed how CFA inactivation suppressed RFA influence on forelimb muscles by obtaining intramuscular electromyographical (EMG) recordings from forelimb muscles during ICMS. EMG recordings showed that despite the presence of evoked movement in CFA, but not RFA, muscle activation in both areas were similarly reduced. These results suggest that the primary reason for the loss of ICMS-evoked movement in RFA is not reduced forelimb muscle activity, but rather a loss of the specific activity between RFA and CFA. Therefore, within the intact motor network of the rat, RFA’s influence on forelimb movement is mediated by CFA, similar to the premotor and motor organization observed in non-human primates.

## INTRODUCTION

It has been well established that a stroke or traumatic brain injury to motor cortex (M1) produces a cascade of secondary histological, physiological, and behavioral events (Nudo 2013). The outcome of injury is compounded as these events initiate and are potentiated by pro- and anti-inflammatory responses, making long-term adaptive or maladaptive plasticity difficult to disentangle from the consequences of the initial injury (Amantea et al., 2009; Donat et al., 2017). As an example, injury induced histological events activated within hours following injury – activation of microglial, inflammatory, and immune response cells; cytokines/chemokines; oxygen/nitrogen free radicals; and several protein-cleaving enzymes – occur contemporaneously with adaptive plasticity, and the reversal of select events have been found to correlate with recovery (Amantea et al. 2009). In effect, since early inflammatory responses can aggravate the initial injury and later responses may be adaptive to recovery mechanisms, these events become entangled with the subsequent long-term neuroplasticity mechanisms that facilitate recovery. Additionally, though it is thought that bi-hemispheric reorganization in spared cortical motor areas are required for motor recovery following injury (Dijkhuizen et al., 2001; Johansen-Berg et al., 2002; Biernaskie et al., 2005; Verley et al., 2018), there is some evidence that the preponderance of functional recovery requires, and may reside within, the ipsilesional hemisphere (Shanina et al., 2006, Harris et al., 2013). Here, intact motor areas extensively reorganize, and compensate over time for the initial M1 injury (Brown et al., 2007; Fridman et al., 2004).

To better understand how the initial injury and not the secondary events influence functional reorganization in forelimb motor areas, studies have examined motor injury through reversable inactivation that preserves cortical tissue and reduces the number of secondary deficits that may influence behavioral outcomes (Matsumura et al., 1991; Kurata & Hoffman, 1994; Kubota, 1996; Martin, 1991, 1993, 1999; Schieber & Poliakov, 1998; Brochier et al., 2004; Sindhurakar et al., 2018), neural activity (Matsumura et al., 1992; Lu & Ashe, 2005; Ohbayashi et al., 2016), evoked muscle responses (Shimazu et al., 2004; Schmidlin et al., 2008), and evoked forelimb movements (Okabe et al., 2016; Dancause et al., 2015; Barry et al., 2014). Using the GABA agonist, muscimol (MUS), we wanted to look at the acute effects of the intact ipsilesional motor network after selective inhibition of rodent M1 (caudal forelimb motor area, CFA) to test the hypothesis that the rodent premotor cortex (rostral forelimb area, RFA) requires an intact CFA to drive forelimb muscle activity. This hypothesis would predict that RFA will be unable to evoke muscle activity to produce forelimb movement in the absence of normal CFA activity. Similar to ventral premotor cortex (PMv) and M1 in primates, the net result would be a motor system where RFA requires an intact CFA to evoke movement (Schmidlin et al., 2008) despite connectivity differences in the proportion of corticofugal fibers between the RFA and CFA of rodents and the PMv and M1 of primates (Rouiller et al., 1993). We found that while some rostral CFA could still evoke movement post-MUS, despite being within the range of MUS spread, the majority of RFA sites observed were unable to evoke forelimb movements, despite being beyond the range of MUS spread. Paradoxically, we found that though RFA forelimb movements were abolished, muscle activation in RFA observed by EMG was similar to CFA sites where forelimb movements could be evoked. The results here are consistent with other studies that show the influence of RFA on forelimb movement is mediated through CFA, and further increases our understanding of consequences of lost motor activity on ipsilesional premotor areas in the absence of secondary events. Our results would support a model where intact premotor circuitry underlies motor recovery of rodent forelimb.

## MATERIALS AND METHODS

### Animals and Environment

N = 24 adult, male Long-Evans hooded rats (3-5 months, weight 350-500g; Envigo, United Kingdom) were procured at 2.5 months of age. Rats were housed in a transparent cage with food and water provided ad-libitum. All rat cages were located in a room with a 12hr:12hr (light:dark) photoperiod and a constant ambient temperature of 22°C. We allowed at least 48hr for rats to acclimate to their housing before experiments were conducted. All daily care procedures and experimental manipulations were reviewed and approved by the University of Kansas Medical Center Institutional Animal Care and Use Committee and adhered to the *Guide for the Care and Use of Laboratory Animals* (Institute for Laboratory Animal Research, National Research Council, Washington, DC: National Academy Press, 1996).

### Experimental Groups and General Procedures

For each rat, anesthesia was induced with an initial administration of isoflurane followed by an injection of ketamine (80-100mg/kg; intra-peritoneal) and Xylazine (5-10mg/kg; intra-muscular). Anesthesia was maintained throughout the procedure with repeated injections of ketamine (10-100 mg/kg/hr ip or im, as needed). To ensure consistent baseline muscle responses that could be influenced by the default positions of the rodent muscles (Sanes et al., 1992), rats were placed in the stereotax with their head and forelimb raised so that forelimbs were suspended with no contact to the surgical bed and no pressure on the forelimbs from the rat’s trunk. A midline incision was made to expose the skull and neck muscles. A second incision was made into the neck muscles above the cervical spine, where an incision was made to the release cerebrospinal fluid within the cisterna magna, reducing intracranial pressure and preventing brain swelling. A craniotomy and dura resection were made across one hemisphere, extending approximately from 5mm rostral and 5mm caudal from rat bregma. Silicon oil was added to the opening to insulate the cortex from the external environment.

Rats were assigned to four different experimental groups. One experiment used neural recordings to establish that our muscimol dosage was localized to CFA and did not spread to RFA (N = 3). A second experimental group was used to determine the consistency of observed forelimb responses in CFA and RFA sites following MUS injection over time (N = 9). Here, following muscimol injection a small number of sites were selected in CFA and RFA and stimulated every 30 minutes over 2 hours to observe perceived changes in current thresholds needed to evoke forelimb responses for each site. Note that because current thresholds for evoked forelimb responses were averaged across all rats, a possibility exists that individual rats with more responsive sites post-MUS injection might bias results relative to rats with less responsive sites.

Once we established the validity of our muscimol manipulation, we obtained a high-density motor map via intracortical microstimulation (ICMS, N = 7). Based on the observed findings from this ICMS group, we concluded that a separate experimental group focusing on electromyography (EMG) recordings was needed, so a set of rats with a lower-density map ICMS group were used to allow sufficient time to record electromyography responses (N = 6).

#### Intracortical Microstimulation

A microelectrode made from a glass micropipette tapered to a fine tip (15-25µm diameter), and filled with 3.5M NaCl, was used for electrical stimulation. Standard ICMS parameters consisted of a 40ms train of 13 200µs monophasic cathodal pulses delivered at 350 Hz from an electrically isolated, constant current stimulator (Nudo et al., 1992, 1996). Pulse trains were delivered at 1HZ intervals. The microelectrode was positioned over the cortical surface containing rat premotor (rostral forelimb area, RFA) and motor (caudal forelimb area, CFA) cortices, and dropped 1650µm from the cortical surface (layer 5). The intensity of the applied current was gradually increased until muscle movement could be identified unambiguously. For this experiment our maximum applied current did not exceed 80µA. We divided observed movements into five categories: distal movements (including wrist and digit extension), proximal movements (elbow flexion), face (including jaw and vibrissae), trunk (including neck and shoulder retraction), and no response. ICMS was delivered to sequential sites, and evoked forelimb motor representations were defined until motor areas were bordered by sites evoking neck, face, or no response. RFA and CFA were typically separated by neck and face representations. For experimental groups, high-and low-density ICMS interpenetration distances were 350µm and 500µm, respectively.

#### Muscimol Inactivation

One microliter of muscimol (MUS, 0.1ng/µl, Sigma-Aldrich) was injected into the approximate center of CFA as determined by the completed ICMS map. Using a 1µl Hamilton microsyringe (Hamilton, Reno, NV, USA) attached to a UMP3 microsyringe pump with Micro4 controller (World Precision Instruments), four boluses of MUS (0.25 µl each, 1µl total) were delivered at a rate of 4nl/sec at depths of 1750µm (1 bolus), 1500µm (2 boli) and 1250µm (1 bolus) from the dorsal surface of CFA. To minimize spilling of the drug outside the insertion tract, the first three injections were followed by 3-minute wait times, and a 5-minute wait time after the 4^th^ injection. Post-MUS ICMS and EMG recording were assessed a minimum of 30-minute post injection, allowing sufficient time for maximal spread. Care was taken to ensure that the distance between the caudal borders of RFA and the MUS injection site exceeded the established maximum radial spread of MUS (**Figure 1**; Martin, 1991).

**Figure 1:**
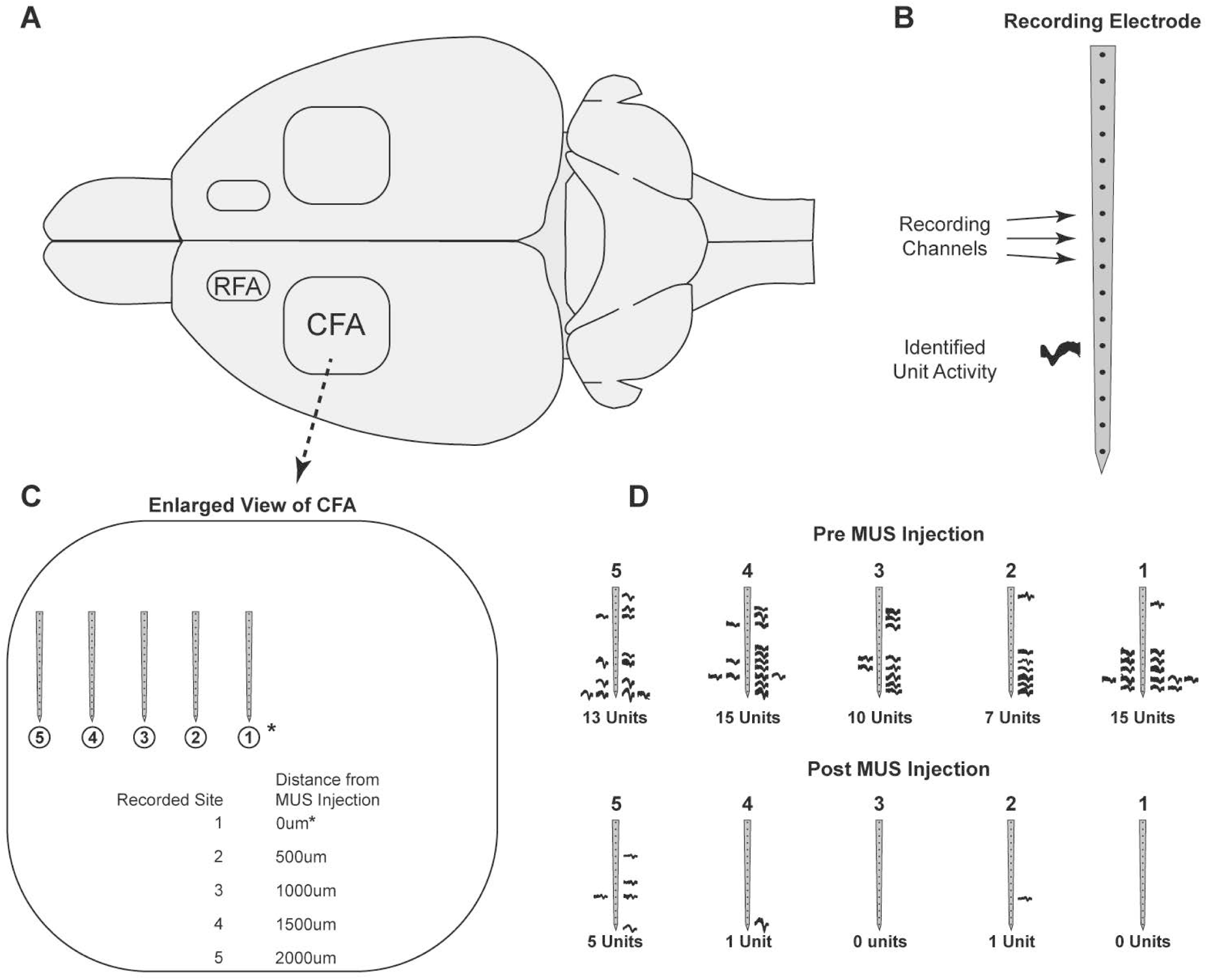
Recorded multi-unit activity in CFA following inactivation by muscimol (MUS). **(A)** Top-down schematic of a rat brain illustrates RFA and CFA in both hemispheres. **(B)** Illustration of 1×16 channel electrode used to acquire neural activity within CFA. Each dot represents a single channel, and a representative waveform for an identified single unit is shown near its respective channel. **(C)** Acquired data was taken at five different recording sites (1-5), each separated by 500µm. 1.0µl MUS was injected at site 1 (represented by the * symbol). After sufficient time for spread (30 minutes), each site was recorded for 5 minutes. **(D)** The total number of identified active units from each electrode at each recording site before and after MUS injection shows that MUS abolished multi-unit activity ∼1500µm from the injection site, and reduced multi-unit activity 2000µm away, inactivating most of CFA.

#### Electromyography (EMG) Electrode Implantation

Pairs of insulated, multi-stranded stainless-steel wires (Type AS 631, Cooner Wire, Chatsworth, CA) were used as intramuscular electrodes. Each wire tip was stripped of 1mm of insulation and folded back on itself into a ‘hook’ electrode. The forelimb skin was shaved, and implantation locations were determined by surface palpation of skin and underlying musculature. EMG electrodes were implanted in four forelimb muscles: biceps brachii, triceps brachii, extensor digitorum communis, and palmaris longus. Due to the small size of rat forelimb muscles, minor variations in placement were to be expected, therefore EMG responses were classified according to functional extensors or flexors movements for wrist or elbow (Liang et al., 1993). EMG electrodes were inserted into the belly of each muscle with a 22-gauge hypodermic needle. Each EMG pair was separated by approximately 5mm and represented one channel.

To verify that the EMG electrodes were within the belly of the muscle, a stimulus isolator (BAK Electronics Inc., Umatilla, FL) was used to deliver a biphasic (cathodic-leading) square-wave current pulse to the muscle through the implanted EMG electrodes. In addition, the impedance between the EMG electrodes was tested via an electrode impedance tester (BAK Electronics, Inc., Umatilla, FL). We tested the impedance values for each EMG electrode relative to: 1) a common ground placed into the base of the tail; and; 2) the other electrode within each muscle pair/channel. EMG electrodes were only accepted if the impedance was ≤10kΩ, direct current delivery to the muscle resulted in contraction of the desired muscle, and the difference in movement threshold between each EMG electrode in the pair/channel was ≤5mA. The external portion of the wires was secured to the skin with surgical glue (3M Vetbond Tissue Adhesive, St. Paul, MN) and adhesive tape.

#### EMG Data Collection

Following ICMS mapping, a 1×16 channel stimulating probe (Neuronexus Technologies, USA) was inserted into select sites in CFA and RFA. Sites were chosen to span the spatial extent for each region and showed comparatively low-to-average ICMS current threshold values. EMG movements were evoked using three 200µs monophasic cathodal pulses delivered at 300Hz from a stimulator (A.M.P.I., Jerusalem, Israel) using a custom-made script (Tucker Davis Technologies, Alachua FI). Stimulus trains were separated by 1s. A 3-pulse train was used as it provided a good signal and avoided potential noise overlap or movement artifact from higher pulse trains and recorded EMG signal, and also has been established previously by our lab (Borrell et al., 2017). Three-pulse stimulation did not induce ICMS movements in our rats, which removed the possibility of movement-related artifact. Pre-MUS injection threshold values were used to evoke EMG movement post-MUS injection. To verify that MUS responses abolished activity, or instead increased threshold to activation, we also applied a separate set of maximum threshold stimulation (80µA) at each site. EMG activity was recorded per site per stimulation threshold for one minute.

#### EMG Data Acquisition

EMG signals from each probe stimulation were recorded and data generated using custom software (Matlab; Mathworks, Inc., Natick, Massachusetts, United States). Each recorded trace from all forelimb muscles was rectified to determine muscle activation (Hudson et al., 2015; Borrell et al., 2017). Stimulus-triggered averages (StTAs) for each recording site for each muscle were also generated, and included in their computation all instances where muscles were both activated and non-activated. For each of the muscles, EMG data were recorded for at least 60 stimulus trains at movement threshold to obtain StTAs and averaged over a set time window of 220ms. StTAs were aligned with the time of the first stimulus (i.e. – 0ms) and included data from −20.2 to +199.8 ms relative to the time of the first stimulus. A muscle was considered active when the average rectified EMG reached a peak ≥2.25 SD above the baseline values in the interval from −20.2 to 0ms and had a total duration of ≥3ms. The stimulus artifact was minimal to absent in EMG recordings with no muscle activation. If an artifact was observed, the amplitude of the artifact was minimal compared to the amplitude of the evoked movement; however, the averaging of the StTA of EMG recordings largely eliminated this type of stimulus artifact. As a result, the stimulus artifact was determined to have minimal to no effect on the recordings. ICMS-evoked EMG potentials were high- and low-pass filtered (30Hz–2.5 kHz), amplified 200–1000 fold, digitized at 5kHz, rectified and recorded on an RX-8 multi-channel processor (Tucker-Davis Technology, Alachua FI). While a separate 60Hz notch filter was not used, 60Hz noise was not evident in the individual traces, and any low-level 60Hz signals were effectively cancelled via signal averaging. EMG signals were stored and analyzed offline.

#### Data Analysis

All between- or within-group difference measures were analyzed using unmatched and matched T-tests, ANOVAs, and appropriate post-hoc tests (GraphPad Prism 5, GraphPad Software, Inc.; JMP 11, SAS, Cary NC). T-tests were used to calculate simple pre and post differences between groups, and one-way and mixed model ANOVAs with Dunnett’s multiple comparison’s test were used for measures with one or two independent variable(s), respectively. The significance threshold was set at p < 0.05.

## RESULTS

### Confirmation of MUS inhibition of CFA unit activity

To verify that the area of functional inactivation coincided with prior histological estimates of MUS spread (Martin, 1991), we tracked multi-unit extracellular activity with a 1×16 channel recording probe (Neuronexus Technologies, USA) within several CFA sites at – and extending from – the injection site, as shown in a representative example in **Figure 1**. A dorsal schematic of the rat brain with locations of motor areas and the recording electrode used to record multi-unit extracellular neural activity are depicted in A and B. Prior to and 30 minutes following injection of MUS into CFA, the electrode recorded neural activity at several sites in 500µm increments from the injection site for 5-minute recording blocks (C). Before inactivation, action potentials were observed from several (7-15) identified units across all sites 2000µm out from the targeted injection site (D). Following the 30-minute inactivation period, multi-unit activity was abolished up to 1500µm away from injection site, with normal unit activity appearing 2000µm from the injection site.

### Effects of MUS on ICMS movement thresholds

Given the length of time required for determination of ICMS movements and current thresholds at sequential sites after MUS injection into CFA (in a 350µm resolution), we sought to determine the impact of post-injection time on the profile of observed movements and thresholds. Though the onset of behavioral deficits after MUS injections in motor cortex has been observed as early as 5 minutes (Okabe et al., 2016) and as long as 24 hours (Martin, 1991) post-injection, it was unclear whether the timing of stimulation for each site (i.e. – earlier or later during the mapping procedure) would bias ICMS map characteristics. In a cohort of eight rats we applied ICMS before and in 30 minute intervals following MUS injection into CFA at the same set of CFA and RFA sites. The total time (two hours) corresponded to the approximate maximum time required to map the entirety of CFA and RFA post-MUS injection at a 350µm resolution.

**Figure 2** shows the current thresholds for evoking movements at various sites over time in CFA and RFA for both saline- and MUS-injected rats, as well the percentage of sites where movement could not be evoked (no response sites). For CFA sites, as expected, both saline and MUS groups showed similar thresholds prior to injection. Using a mixed model ANOVA analysis to assess the effect of injection (MUS or saline) and time (pre; post 1,2,3,4) on current thresholds in CFA, we found a significant effect for both time [F_(4,328.2)_ = 17.91; p < 0.0001] and injection*time interaction [F_(4,328.2)_ = 4.99; p < 0.001]. Dunnett’s post hoc tests showed that post-MUS current thresholds were significantly increased relative to pre-MUS thresholds at 30 (p < 0.0001), 60 (p < 0.0001), 90 (p < 0.0001) and 120 (p < 0.0001) minutes post-injection. By comparison, post-saline current thresholds were found to differ relative to pre-saline thresholds at 90 (p < 0.01) and 120 (p < 0.0001) minutes post injection. These results would suggest that the current thresholds of our rats naturally increased as a function of time under Ket/Xyl anesthesia, and that MUS hastens this increase, such that CFA current thresholds are significantly increased 30 minutes post injection. Note that while, by definition, ICMS elicited forelimb movements at all sites prior to MUS or saline injections, trunk/facial movements were sometimes evoked at these forelimb sites after injection. MUS injections also resulted in an increase in the number of no response sites observed in CFA relative to saline injected rats (closed triangle vs closed circles).

**Figure 2:**
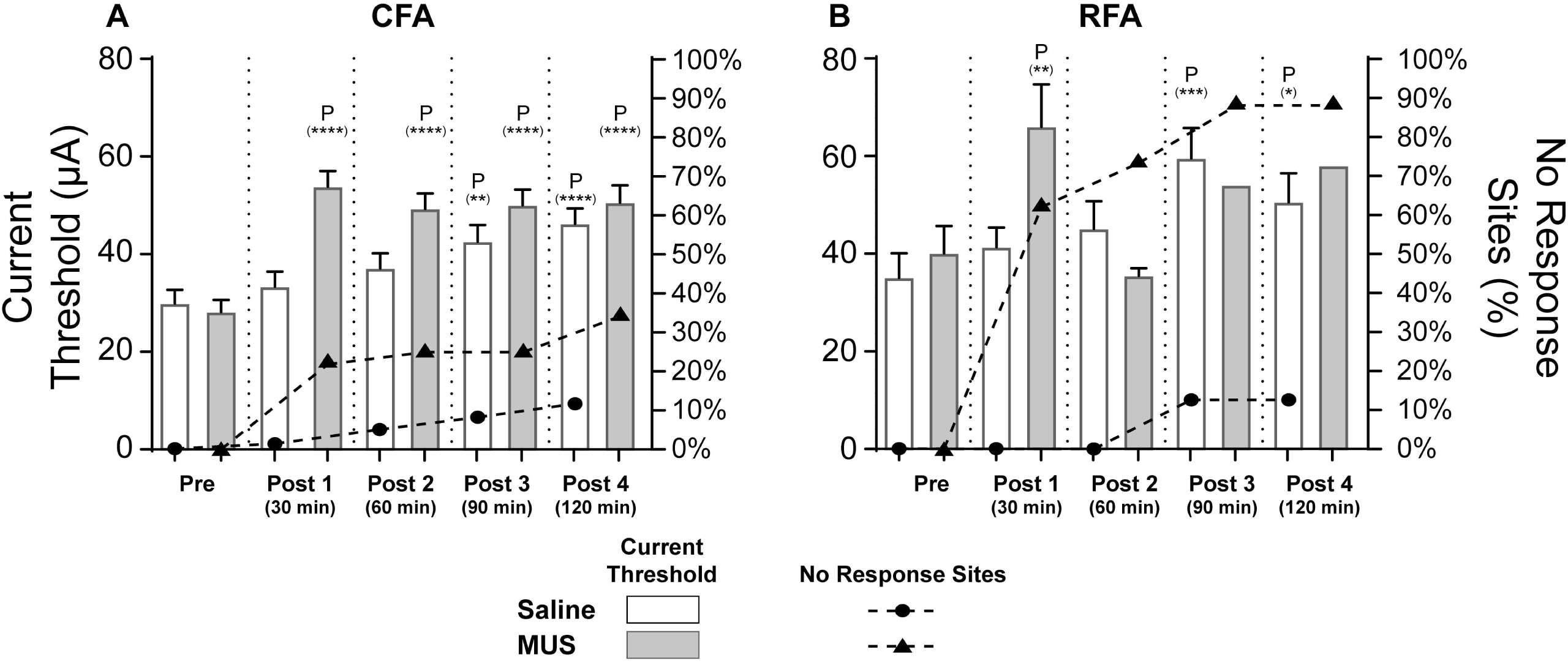
Observed map changes are relatively stable over the acute period following inactivation. Bar and line graphs show the increases in threshold and no response sites as a function of surgical time. White bars represent a (phosphate-buffered) saline-injected control group and gray bars represent MUS-injected group. In CFA, the saline control group shows that the minimum current necessary to activate forelimb movements increases steadily over time until 90 minutes into the experiment: At CFA, mixed model ANOVA analysis using injection (MUS or saline) and time (Pre; Post 1, 2, 3, 4) show a significant effect of both time [F_(4,328.2)_ = 17.91; p < 0.0001] and injection*time interaction [F_(4,328.2)_ = 4.99; p < 0.001]. Dunnett’s post hoc tests show significant differences in both saline and MUS injections: current thresholds were significantly increased 90 min (Post 3, p < 0.01) and 120 min (Post 4, p < 0.001) from pre- to post-saline injection; and, current thresholds were significantly increased at 30 (p < 0.0001), 60 (p < 0.0001), 90 (p < 0.0001) and 120 (p < 0.0001) minutes from pre- to post-MUS injection. In RFA, there is also a significant effect of both time [F_(4,38.79)_ = 4.92; p < 0.01] and injection*time interaction [F_(4,38.79)_ = 2.97; p < 0.05]. Dunnett’s post hoc tests show significant differences in both saline and MUS injections: current thresholds were significantly increased at 90 min (Post 3, p < 0.001) and 120 min (Post 4, p < 0.05) from pre- to post-saline injection; and, current thresholds were significantly increased 30 min (Post 1, p < 0.01) from pre- to post-MUS injection. A small number of sites in CFA and RFA (10%) showed no response in the saline control group by the end of the experiment (closed-circle line graph). By post-MUS injection time 4, 35% of CFA sites no longer evoked forelimb movements. No significant increases in threshold values could be demonstrated in RFA of the MUS group, due to the significant loss of forelimb sites. By RFA post-injection time 4, 9 out of the 10 tested sites no longer evoked forelimb movements (closed-triangle line graph). Note that columns for post-MUS injection times 3 and 4 represent only one active RFA site, and do not have error bars. (P) Significant differences from Pre current threshold. (****) P < 0.0001. (**) P < 0.01. (*) P < 0.05.

Mixed model ANOVA analysis for RFA shows a similar significant effect in both time [F_(4,38.79)_ = 4.92; p < 0.01] and injection*time interaction [F_(4,38.79)_ = 2.97; p < 0.05]. Dunnett’s post hoc tests show a significant increase in RFA current thresholds at 30 minutes after MUS injection (Post 1, p < 0.01). After, ICMS-evoked movements in RFA were largely abolished after MUS injection into CFA, with the majority of sites showing no response to ICMS (closed triangles). Consequently, as ICMS-evoked movements in RFA were largely abolished after MUS injection into CFA, this precluded pre-post threshold comparisons at later time points 60, 90, and 120 minutes post-MUS injection. Like CFA, post-saline injection time points were found to differ 90 (p < 0.001) and 120 (p < 0.05) minutes post injection, supporting the observation that current thresholds naturally increase over time as a function of time under Ket/Xyl anesthesia.

We also observed that while saline-injected rats showed qualitatively consistent forelimb movement responses to ICMS, the ICMS-evoked movements of MUS-injected rats were much less consistent: Evoked movements sometimes switched between the ipsi- and contra-lateral forelimbs, or no response/non-forelimb sites sometimes evoked forelimb movements at later time points, though in the latter case these sites required higher than average currents to evoke movements (>60µA). While the increase in no response sites following MUS injection suggests that CFA function has been partially – but not completely – impaired, the increased movement variation observed across the two hours following MUS injection may reflect a fundamental dysregulation of the CFA- to-forelimb muscle pathway that results in inconsistent activation of forelimb muscles.

### MUS reduces the size of ICMS-evoked forelimb area in CFA and abolishes most evoked forelimb movement in RFA

**Figure 3A** shows a representative dorsal view of a rat brain containing both CFA (white dashed outline) and RFA (red dashed outline) movement representations before and after MUS injection into CFA. Each color represents a different observed ICMS-evoked movement response category. Forelimb movements are indicated in blue (proximal) and green (distal); non-forelimb movements are indicated in pink (trunk) and yellow (face); sites which show no response are indicated in black. MUS injections (dashed circle) in CFA significantly reduced the total percentage of ICMS-evoked forelimb movements and increased no response sites **(Fig. 3B)**. When averaged across all rats, CFA sites that evoked forelimb responses pre-MUS injection largely showed no response post-MUS injection even at maximum current thresholds (53.6%), or evoked trunk (6.6%) or facial (3.9%) movements. As a result, the total area of evoked proximal and distal forelimb movements in CFA was decreased significantly after MUS injections [*proximal*: t_(1, 6)_ = 4.141, p < 0.01; *distal*: t_(1, 6)_ = 3.79, p < 0.01], while the no response area significantly increased [t_(1, 6)_ = 11.22, p < 0.0001] **(Figure 4)**. Like figure 3A implies, we also noted that sites in CFA that evoked distal forelimb movements pre-MUS were more likely to evoke proximal movements post-MUS. That is, post-MUS forelimb movements were likely to be proximal. In RFA, forelimb movements evoked prior to MUS injection were largely abolished post-MUS injection: a few sites could still evoke forelimb movements (8.3% of sites observed), but most sites that evoked forelimb movements pre-MUS showed either no response at maximum current thresholds (73.3%) or evoked trunk/facial movements (18.3%, data not shown). In total, MUS injection into CFA appears to disproportionally attenuate distal responses in CFA and both distal and proximal responses in RFA.

**Figure 3:**
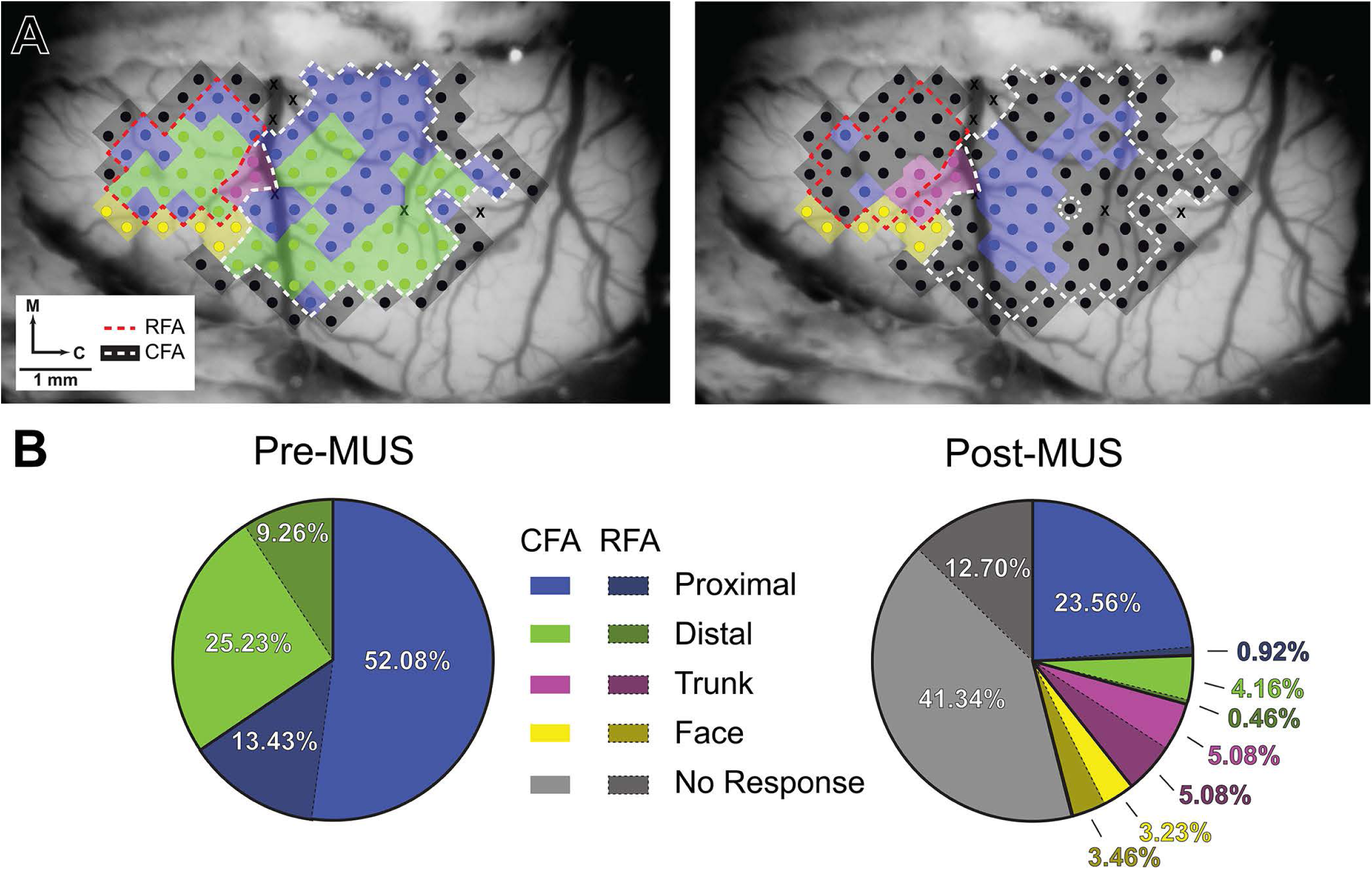
MUS injection into CFA reduces ICMS-evoked forelimb representation in both CFA and RFA. **(A)** Top-down example of an exposed rat brain showing a high-density ICMS map for CFA (white dashed outline) and RFA (red dashed outline) before and after MUS injection (injection site shown here as a small dashed circle in the approximate central location of CFA). Each site containing ICMS-evoked movement is color-coded: blue indicates proximal (elbow flexion), green indicates distal (wrist extension), pink indicates trunk, yellow indicates face/vibrissae, and black/gray indicates sites no response to stimulation even at the experimental maximum applied current (80µA). Sites marked X were skipped over to avoid damage to major blood vessels. In this rat, MUS injected into CFA largely abolished the distal forelimb activity evoked in RFA and the caudal sites in CFA largely showed no response to ICMS. **(B)** Total percentage for each evoked movement in CFA and RFA before and after MUS injection. In the first pie chart, the total percentage of proximal and distal forelimb sites are shown for both CFA (light-colored) and RFA (dark-colored) averaged across N = 7 rats. The second pie chart shows changes in CFA and RFA due to MUS. We see that over half of the total proximal and distal sites have showed no response to ICMS or evoked trunk/facial/vibrissae movements. In RFA, almost the entirety of proximal and distal sites has been abolished across all rats. (M = rostral, C = caudal).

**Figure 4:**
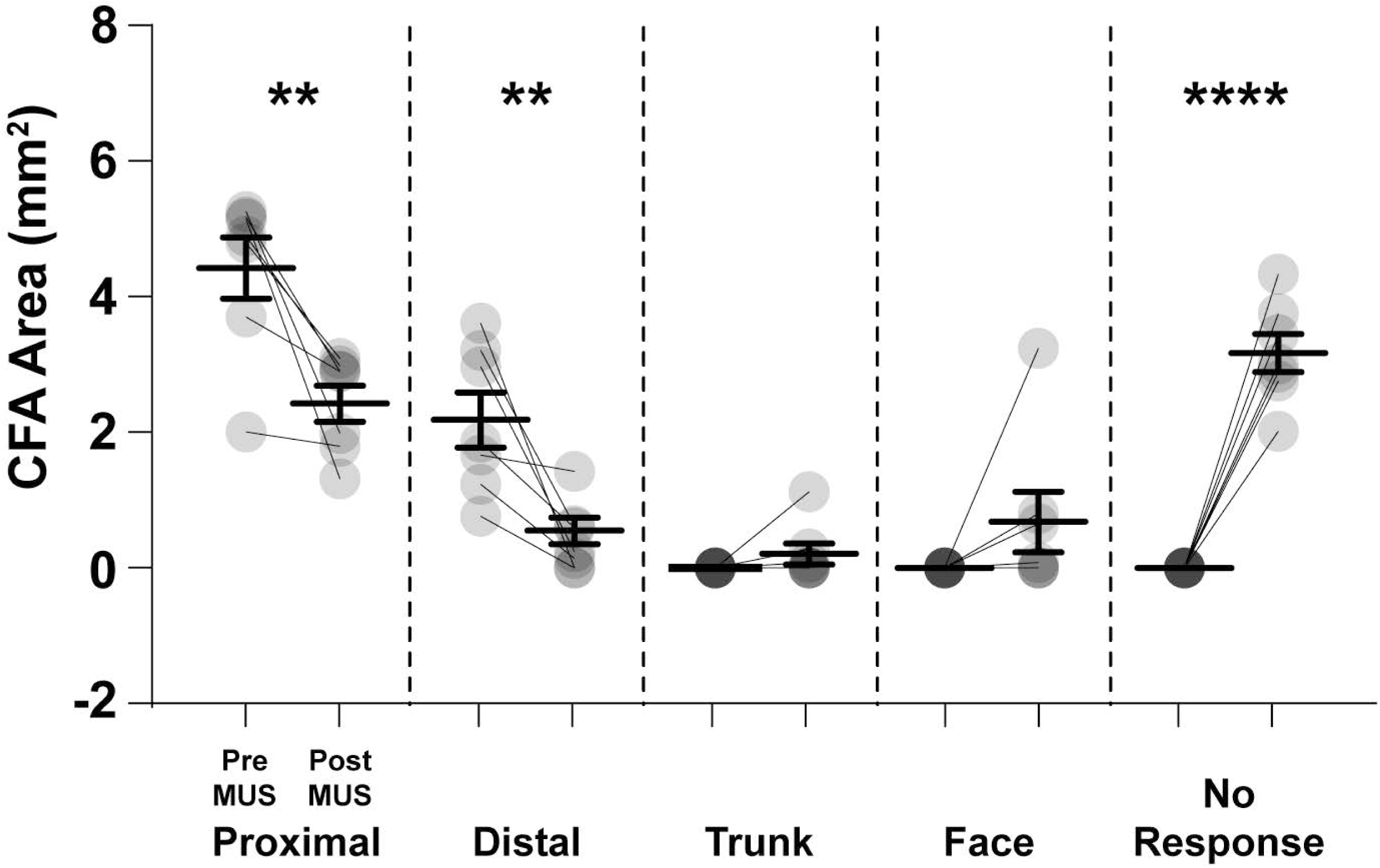
MUS injections significantly reduce area of evoked-forelimb responses in CFA. Column scatter-plots show group mean and individual subject (N = 7) cortical surface area for all ICMS-evoked responses and no response sites in CFA before and 30 minutes after MUS injection. On average, the area of evoked proximal and distal forelimb movements is significantly smaller after MUS injection [*proximal*: t_(1, 6)_ = 4.141, p < 0.01; *distal*: t_(1, 6)_ = 3.79, p < 0.01], with a significant increase in no response sites [t_(1, 6)_ = 11.22, p < 0.0001]. Face and trunk sites did not significantly increase following injection.

**Figure 5A** shows that in CFA sites where responses persisted, the average current threshold required to evoke forelimb movements was significantly increased (but still within a normal physiological range) compared to prior pre-MUS current thresholds [Rat 1: t_(1, 6)_ = 4.868, p < 0.01; Rat 2: t_(1, 12)_ = 3.295, p < 0.01; Rat 3: t_(1, 17)_ = 4.417, p < 0.001; Rat 4: t_(1, 19)_ = 10.05, p < 0.0001; Rat 5: t_(1, 25)_ = 4.131, p < 0.001; Rat 6: t_(1, 26)_ = 8.721, p < 0.0001; Rat 7: t_(1, 17)_ = 12.37, p < 0.0001]. In RFA, the reduced number of responsive forelimb sites observed post-MUS were too few to statistically compare pre- and post-MUS current thresholds, but of the responsive sites present we did note an increase in average current thresholds required to evoke forelimb movements. Of the remaining present forelimb sites in CFA and RFA, the occurrence of distal forelimb movements was reduced: distal forelimb movements were evoked in approximately one-third of CFA and one-half of RFA pre-MUS injection but only one-seventh in CFA and one-fifth in RFA post-MUS. Stimulation of responsive sites largely evoked proximal movements in areas that previously evoked distal movements. Additionally, though post-MUS sites in RFA consistently showed no response, several sites in CFA that showed no response had brief instances where movements were initially evoked across several stimulation trains lasting several seconds, at which point movement immediately ceased and could no longer be evoked at that site. Taken together, we found that acute inactivation of CFA due to MUS injection abolishes forelimb responses in RFA and reduces the total area of CFA representation, predominantly through distal forelimb inactivation and decreases in the capacity to maintain movement responses.

**Figure 5:**
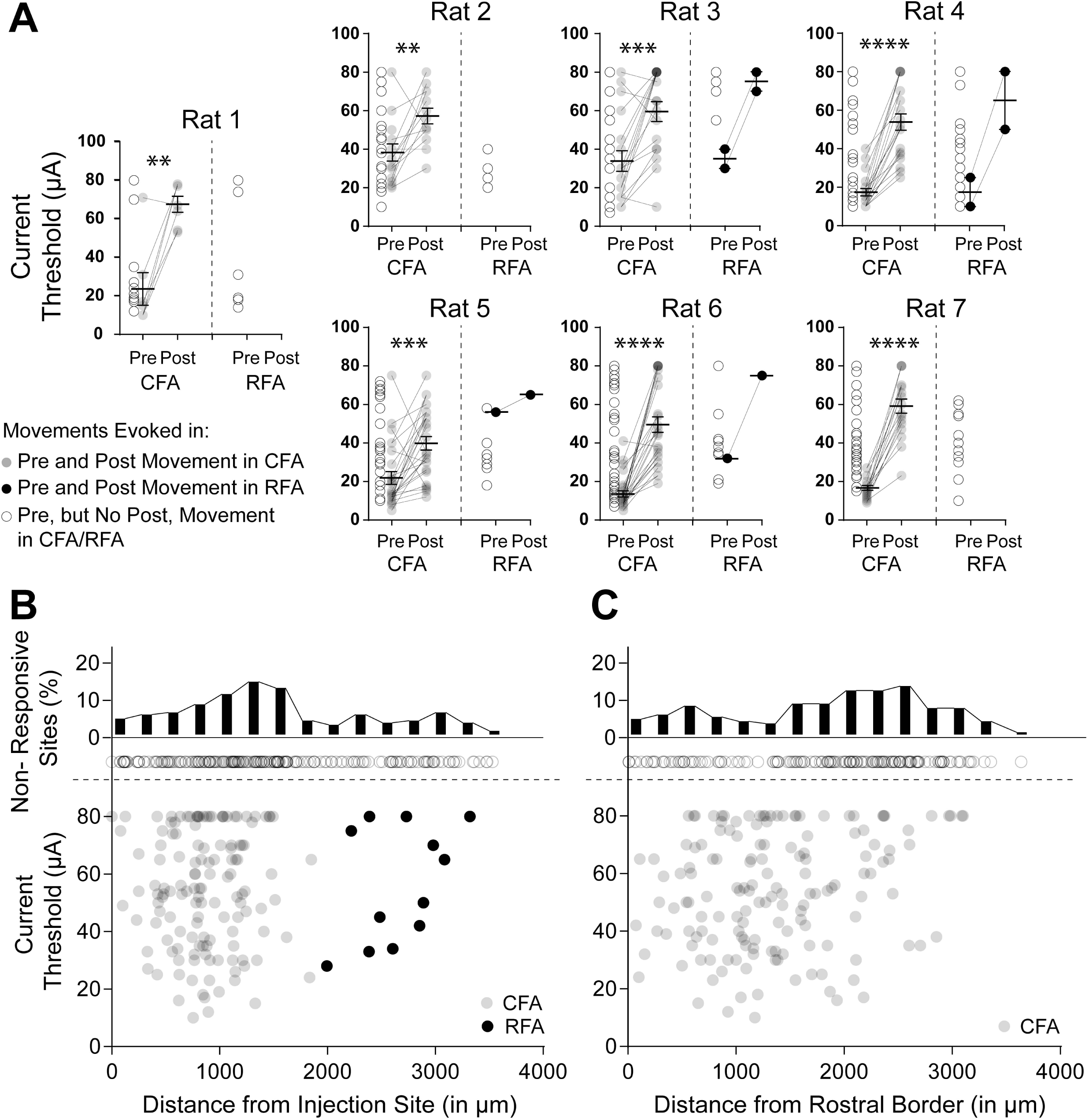
Distribution of no response sites, but not site thresholds, shows rostral-caudal bias. Gray and black circles represent forelimb movements in CFA and RFA that could be evoked both before (pre) and after (post) MUS injection, respectively. Open circles represent pre-injection movements that showed no response to ICMS post-MUS injection. Open circles are located at pre current threshold value in (A) or with respect to their distance from the injection site in (B,C). **(A)** Column scatter-plots show group mean and individual site thresholds for each rat with ICMS maps before and after MUS injection. Within each CFA the average threshold for intact sites was found to significantly increase after MUS injection [Rat 1: t_(1, 6)_ = 4.868, p < 0.01; Rat 2: t_(1, 12)_ = 3.295, p < 0.01; Rat 3: t_(1, 17)_ = 4.417, p < 0.001; Rat 4: t_(1, 19)_ = 10.05, p < 0.0001; Rat 5: t_(1, 25)_ = 4.131, p < 0.001; Rat 6: t_(1, 26)_ = 8.721, p < 0.0001; Rat 7: t_(1, 17)_ = 12.37, p < 0.0001]. While RFA sites were largely abolished and too few remained to perform statistical comparisons, the sites that remained tended to have lower threshold values compared to other pre-injection sites. **(B**,**C)** Scatterplot and histogram reveal that no response sites are not concentrated nearer to injection site, but rather towards the caudal half of CFA. Distance from injection site was compared against both: 1) current threshold values of post-injection forelimb sites in CFA (gray, B and C) and RFA (black, B only), and 2) the percentage of no response sites (open circles, B and C). As shown in B, no notable correlation was found between post CFA and RFA values and the distance from the injection site, suggesting that remaining site activity did not vary as a function of proximity to the MUS injection site. However, when looking at the percentage of responsive sites that showed no response post-injection (binned in 250µm increments), C shows that no response site distribution was biased not toward the injection site (B), but rather towards the caudal half of CFA. Threshold values were not found to correlate with this rostral-caudal bias.

### CFA inactivation produces global increase in thresholds and regional loss of evoked movements

In an attempt to identify a pattern of site activity post-MUS injection, we considered the possibility that the observed movements or lack thereof for each site post-MUS might be related to the distance of each site relative to the MUS injection site. We also observed a potential caudal bias with respect to the no response sites in CFA, and so considered the possibility that observed movements were dependent on distance relative to some rostral-caudal position. To determine if a spatial relationship existed between injection location/rostral border and ICMS-evoked responses, we looked at the distribution of current threshold values relative to the distance from the injection site/rostral border. We defined a rostral border for CFA using non-forelimb sites prior to MUS injection and oriented orthogonal to the sagittal suture skull line. Figures **5B** **and C** compare threshold values for all sites in CFA and RFA that elicited a movement (forelimb, trunk, and neck) post-MUS injection against their distance from the injection site/rostral border. After MUS injection, we found that site threshold values neither varied relative to their distance from the injection site (**Fig. 5B**) nor rostral border of CFA (**Fig. 5C**). When we isolated each movement, we found neither sites that maintained evoked forelimb movements nor transformed into evoked trunk or facial movements correlated with distance. Distance from the injection site appeared to inversely correlate to amount of no response sites. However, this is likely due to the increased area canvased as a function of radial distance to the injection site.

When distance from the rostral border of CFA was assessed, we observed what appeared to be a bimodal distribution of no response sites in the rostral and caudal halves of CFA, with a clear greater concentration of no response sites towards the caudal half. Together, figures B and C show that while no response sites are *not* concentrated near the injection site, they are not stochastically distributed throughout the areal extent of CFA, instead having a higher likelihood of being caudally distributed. As injections were given at the approximate center of the CFA ICMS maps, these data might suggest a heretofore unknown division in GABAergic or glutamatergic sensitivity or functional subdivisions within CFA.

### EMG activity patterns in the rat

Prior to testing the effects of MUS on ICMS-evoked EMG responses, we wanted to establish both the normal patterns of activation as a byproduct of our experimental parameters and ensure that the EMG features were consistent with what should be expected given the observed proximal and distal forelimb movements. Stimulation of either proximal or distal sites by 3-pulse ICMS trains were found to activate predominantly biceps, wrist extensors, and flexors. Triceps activity was minimal, with comparably flat peaks. Stimulus-triggered averages were generated using the EMG data output from at least 60 3-pulse ICMS trains (StTAs, See Methods). Multivariate analyses comparing the EMG peak amplitude during fixed 3-pulse ICMS show a strong positive correlation between biceps and wrist flexors (r = 0.61, p < 0.0001), and a moderate positive correlation between wrist extensors and the more minimally active triceps (r = 0.40, p = 0.01; **Figure 6**). That is, instances where wrist flexors or triceps are maximally activated are likely to occur when biceps or wrist extensors are maximally active, respectively. These correlations, and the lack of any notable correlation between the StTA peak amplitudes for other muscle pairs suggest that the forelimb movements that activate biceps (or wrist extensors) recruit wrist flexors (or triceps) in a similar manner. Alternatively, some of the observed wrist flexor (or extensor) activation could be due to artifact from bicep (or triceps) activation. The later possibility is unlikely as our 3-pulse stimulation did not produce movement, and is also not supported by the low latency correlations between the StTA activation of biceps to wrist flexors and of triceps to wrist extensors. Instead, we find strong positive correlations between the timing of the more active bicep and wrist extensors (r = 0.62, p < 0.0001), and between the less active wrist flexors and triceps (r = 0.40, p = 0.01). The dual correlations within peak amplitude and latency are expected given the separate proximal and distal forelimb movements: proximal bicep flexion would predominantly activate biceps, with potential secondary sactivation of wrist flexors; and distal wrist extension would predominantly activate wrist extensors with potential activation of triceps to stabilize the upper forelimb during wrist extension.

**Figure 6:**
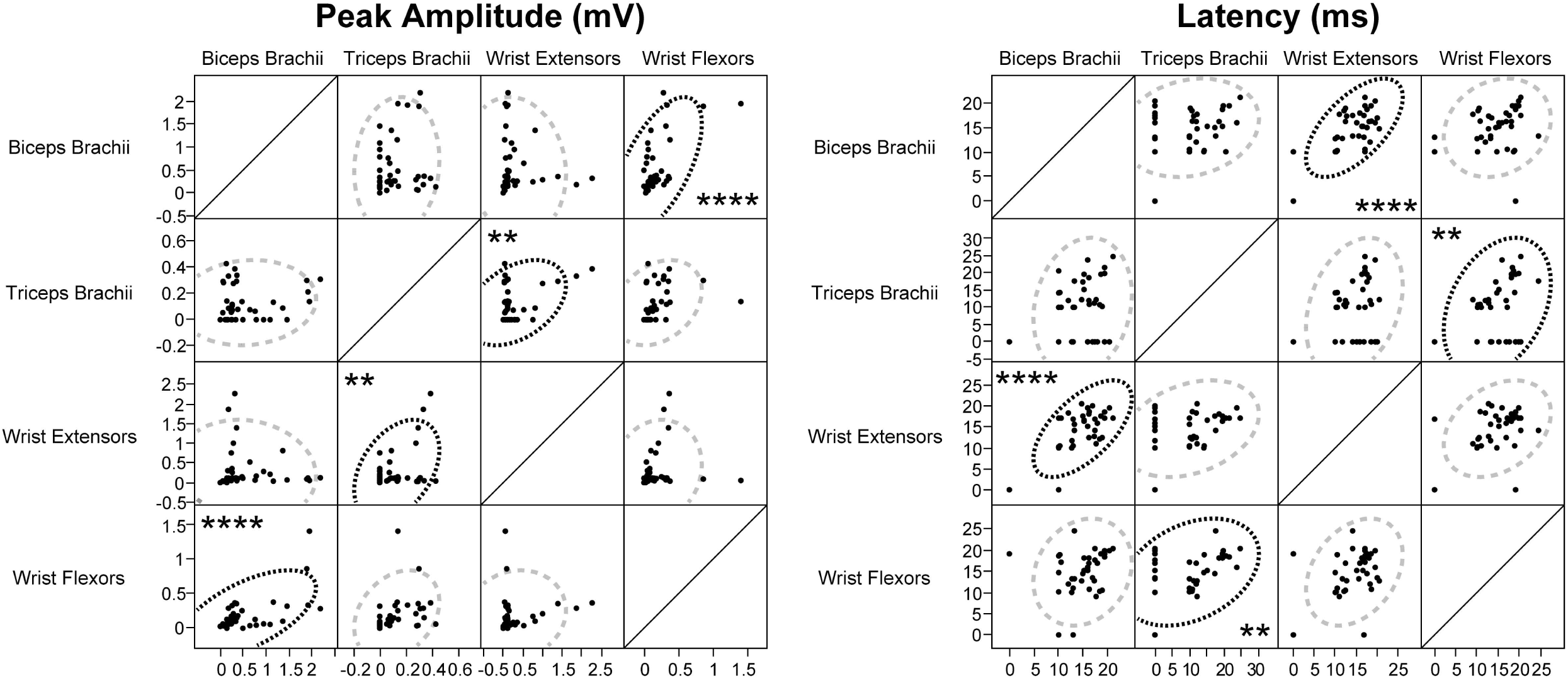
The pattern of amplitude and timing of forelimb muscle activity during cortical stimulation reflect the dual distal and proximal movements. Correlation matrices show the stimulus-triggered average (StTA) amplitude and latency values for EMGs recorded from all four muscles. Significant correlations are highlighted with black dotted lines. We observed strong correlations in biceps / wrist flexors (r = 0.61, p < 0.0001) and moderate correlations in wrist extensors / triceps (r = 0.40, p < 0.01) for StTA peak amplitude. Likewise, we observed strong correlations in biceps / wrist extensor (r = 0.62, p < 0.0001) and moderate correlations in triceps and wrist flexors (r = 0.40, p < 0.01) for StTA latency.

We had an *a priori* expectation that larger elbow flexion (proximal) movements would elicit greater activation amplitude measures relative to the smaller wrist extension (distal) movements, and could be used as a clear demarcation for movement type. Unsurprisingly, bicep activation had significantly higher amplitude in elbow flexion relative to wrist extension movements [F_(1, 9.228)_ = 17.26; p < 0.01]. What was surprising, however, was that the relative responses from the other three muscles did not significantly differ between proximal or distal site stimulation: biceps are the only muscle group that significantly differs in peak amplitude during activation as a function of movement type. These and the above patterns help to provide an initial baseline prior to inactivation and could also be used to assess the reliability of our experimental parameters as a tool to measure changes after inactivation.

We also looked to see whether notable differences in the amplitude or the timing of muscle activation existed between CFA and RFA before MUS injection. We observed significant difference in latency of muscle activation in biceps, which were on average 2.4ms faster in activation in CFA relative to RFA. For peak amplitude, we also found that wrist extensors show greater amplitude when stimulation occurs in RFA, but biceps show greater amplitude for CFA stimulation. While these results may be a consequence of our dataset, it may also suggest a bias of bicep muscle activation with respect to CFA, and wrist extensor activation with respect to RFA.

### Inactivation of CFA suppresses baseline and evoked forelimb muscle activity

Our next experiment addressed how CFA inactivation indirectly abolished RFA evoked movements yet leaves some CFA movement intact. One possibility is that MUS injections into CFA block the indirect RFA-to-motor neuron connectivity that routes through CFA. CFA activity would be largely preserved due to the direct and artificial ICMS of descending CFA to motor neuron fibers, but the natural neuronal firing from RFA would be insufficient to drive CFA activity (i.e. – a loss of activation). Alternatively, perhaps RFA activity “makes it through” to drive muscle activation, but a required pattern or summation of muscle activity coordinated by CFA is necessary for movement to occur (i.e. – a loss of timing). To test one of these possibilities, we looked at EMG activity in forelimb muscles at select proximal or distal sites in CFA and RFA before and after MUS injection. **Figure 7** shows a representative example of MUS suppression effects on the StTAs for all four muscles groups at proximal and distal sites in CFA and RFA. For each of the four areas given 3-pulse ICMS, each pre current threshold value was applied before (in grey) and after (in red) injection. Vertical dashed lines represent timing of stimulation. Biceps and wrist flexors were predominantly activated during proximal movement, with no notable activity in either wrist extensors and triceps. Likewise, wrist extensors and some biceps were predominantly activated at distal movement sites in both CFA and RFA, with minimal to no activity in wrist flexors or triceps. This was observed in most instances where pre-MUS threshold values were used in CFA and RFA. For these examples, peak amplitude and mean baseline for evoked-EMG measures observed in all muscles were attenuated following MUS injection. **Figure 8** shows that when threshold values (white boxes) were used across all rats, we found significant reductions in the peak amplitude of biceps, wrist flexors, and wrist extensors (**See Table 1**). Mean baseline amplitude for biceps, wrist extensors, wrist flexors, and triceps during stimulation trains were also reduced, supporting the conclusion that, although some of CFA can still evoke forelimb movement, forelimb muscle activation in CFA has been impaired as a function of direct MUS injection.

**Table 1:**
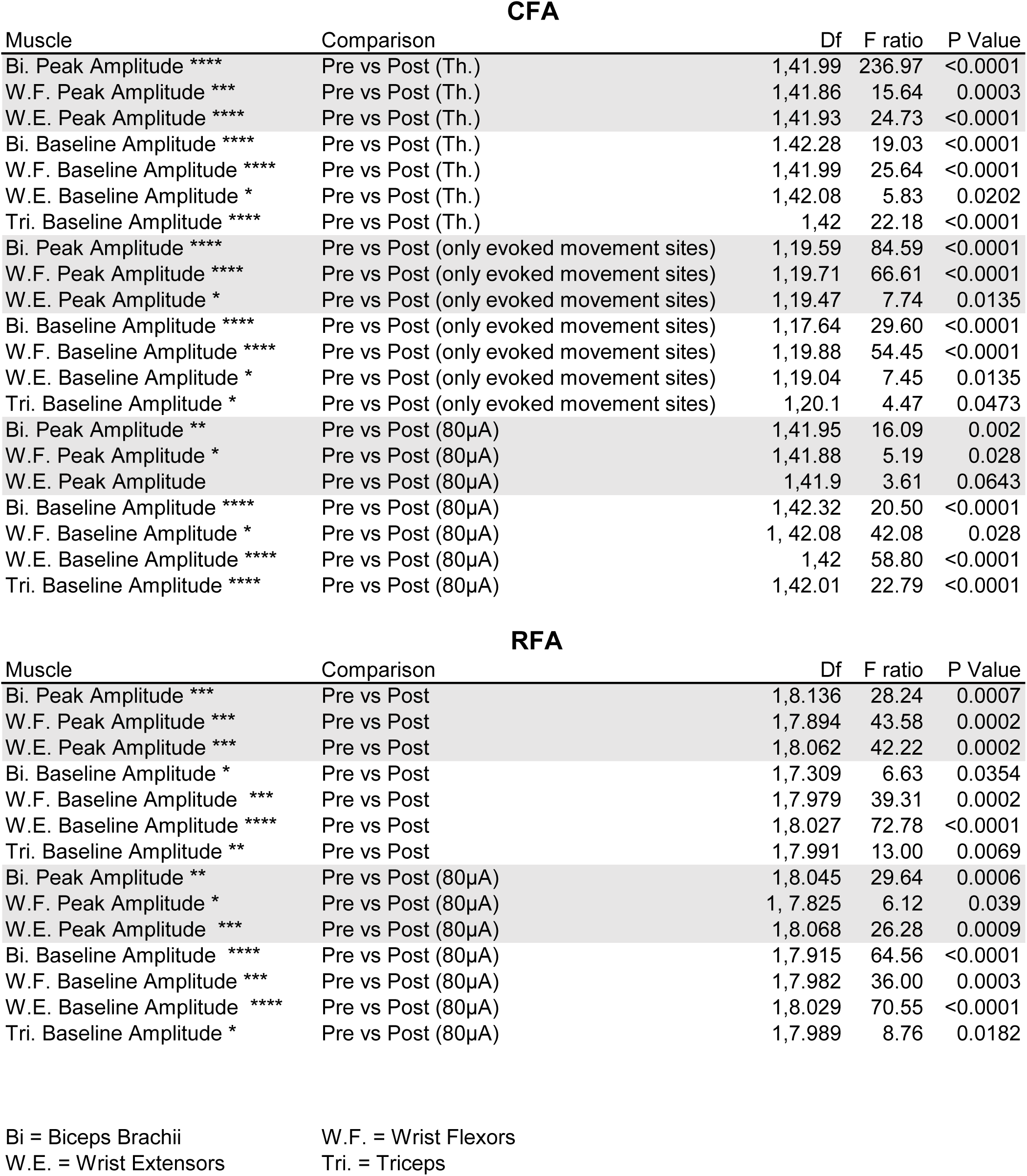
Statistical summary of StTA comparisons made in Figure 8.

| Muscle | Comparison | Df | F ratio | P Value |
| --- | --- | --- | --- | --- |
| Bi. Peak Amplitude **** | Pre vs Post (Th.) | 1,41.99 | 236.97 | <0.0001 |
| W.F. Peak Amplitude *** | Pre vs Post (Th.) | 1,41.86 | 15.64 | 0.0003 |
| W.E. Peak Amplitude **** | Pre vs Post (Th.) | 1,41.93 | 24.73 | <0.0001 |
| Bi. Baseline Amplitude **** | Pre vs Post (Th.) | 1,42.28 | 19.03 | <0.0001 |
| W.F. Baseline Amplitude **** | Pre vs Post (Th.) | 1,41.99 | 25.64 | <0.0001 |
| W.E. Baseline Amplitude * | Pre vs Post (Th.) | 1,42.08 | 5.83 | 0.0202 |
| Tri. Baseline Amplitude **** | Pre vs Post (Th.) | 1,42 | 22.18 | <0.0001 |
| Bi. Peak Amplitude **** | Pre vs Post (only evoked movement sites) | 1,19.59 | 84.59 | <0.0001 |
| W.F. Peak Amplitude **** | Pre vs Post (only evoked movement sites) | 1,19.71 | 66.61 | <0.0001 |
| W.E. Peak Amplitude * | Pre vs Post (only evoked movement sites) | 1,19.47 | 7.74 | 0.0135 |
| Bi. Baseline Amplitude **** | Pre vs Post (only evoked movement sites) | 1,17.64 | 29.60 | <0.0001 |
| W.F. Baseline Amplitude **** | Pre vs Post (only evoked movement sites) | 1,19.88 | 54.45 | <0.0001 |
| W.E. Baseline Amplitude * | Pre vs Post (only evoked movement sites) | 1,19.04 | 7.45 | 0.0135 |
| Tri. Baseline Amplitude * | Pre vs Post (only evoked movement sites) | 1,20.1 | 4.47 | 0.0473 |
| Bi. Peak Amplitude ** | Pre vs Post (80µA) | 1,41.95 | 16.09 | 0.002 |
| W.F. Peak Amplitude * | Pre vs Post (80µA) | 1,41.88 | 5.19 | 0.028 |
| W.E. Peak Amplitude | Pre vs Post (80µA) | 1,41.9 | 3.61 | 0.0643 |
| Bi. Baseline Amplitude **** | Pre vs Post (80µA) | 1,42.32 | 20.50 | <0.0001 |
| W.F. Baseline Amplitude * | Pre vs Post (80µA) | 1, 42.08 | 42.08 | 0.028 |
| W.E. Baseline Amplitude **** | Pre vs Post (80µA) | 1,42 | 58.80 | <0.0001 |
| Tri. Baseline Amplitude **** | Pre vs Post (80µA) | 1,42.01 | 22.79 | <0.0001 |

RFA
| Muscle | Comparison | Df | F ratio | P Value |
| --- | --- | --- | --- | --- |
| Bi. Peak Amplitude *** | Pre vs Post | 1,8.136 | 28.24 | 0.0007 |
| W.F. Peak Amplitude *** | Pre vs Post | 1,7.894 | 43.58 | 0.0002 |
| W.E. Peak Amplitude *** | Pre vs Post | 1,8.062 | 42.22 | 0.0002 |
| Bi. Baseline Amplitude * | Pre vs Post | 1,7.309 | 6.63 | 0.0354 |
| W.F. Baseline Amplitude *** | Pre vs Post | 1,7.979 | 39.31 | 0.0002 |
| W.E. Baseline Amplitude **** | Pre vs Post | 1,8.027 | 72.78 | <0.0001 |
| Tri. Baseline Amplitude ** | Pre vs Post | 1,7.991 | 13.00 | 0.0069 |
| Bi. Peak Amplitude ** | Pre vs Post (80µA) | 1,8.045 | 29.64 | 0.0006 |
| W.F. Peak Amplitude * | Pre vs Post (80µA) | 1, 7.825 | 6.12 | 0.039 |
| W.E. Peak Amplitude *** | Pre vs Post (80µA) | 1,8.068 | 26.28 | 0.0009 |
| Bi. Baseline Amplitude **** | Pre vs Post (80µA) | 1,7.915 | 64.56 | <0.0001 |
| W.F. Baseline Amplitude *** | Pre vs Post (80µA) | 1,7.982 | 36.00 | 0.0003 |
| W.E. Baseline Amplitude **** | Pre vs Post (80µA) | 1,8.029 | 70.55 | <0.0001 |
| Tri. Baseline Amplitude * | Pre vs Post (80µA) | 1,7.989 | 8.76 | 0.0182 |

Bi = Biceps Brachii W.E. = Wrist Extensors
W.F. = Wrist Flexors Tri. = Triceps

**Figure 7:**
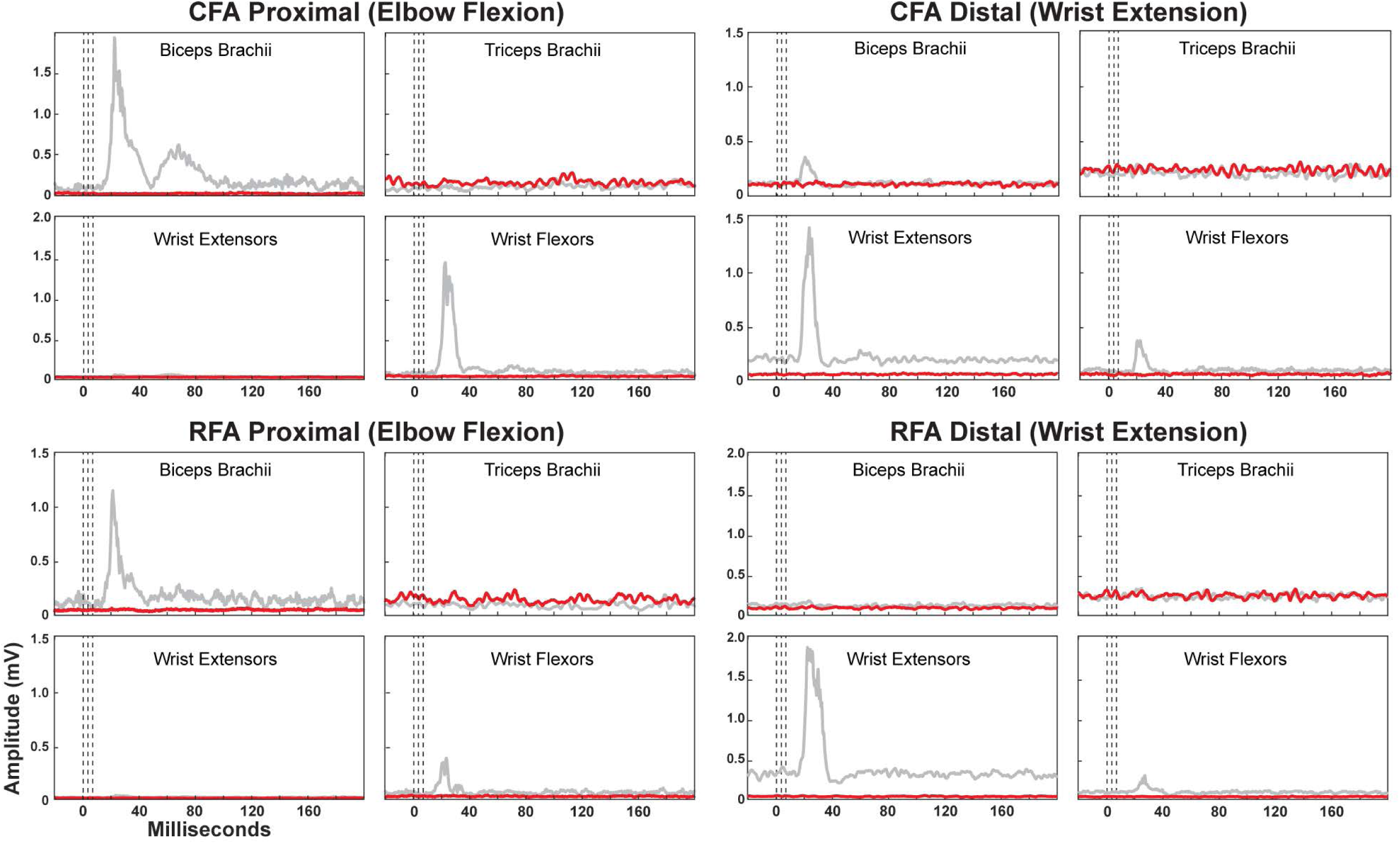
MUS inactivation of CFA reduces EMG forelimb properties. Representative example of StTAs gathered from EMG activity measured at each forelimb muscle before (gray lines) and after (red lines) MUS injection into CFA. Here, distal and proximal sites from CFA and RFA identified by 13-pusle ICMS were chosen, and StTAs were obtained using 3-pulse stimulation (dotted lines). In both the CFA and RFA sites that produced proximal forelimb movements there is prominent activation of biceps and some wrist flexors (note different y-axis values). Following injection both the peak amplitude and mean baseline values for all active muscles were reduced. Likewise, in CFA and RFA sites that evoked distal forelimb movement, we observed prominent wrist extensor activation and some biceps and wrist flexors prior to MUS inhibition, and these activations were similarly attenuated following MUS inactivation.

**Figure 8:**
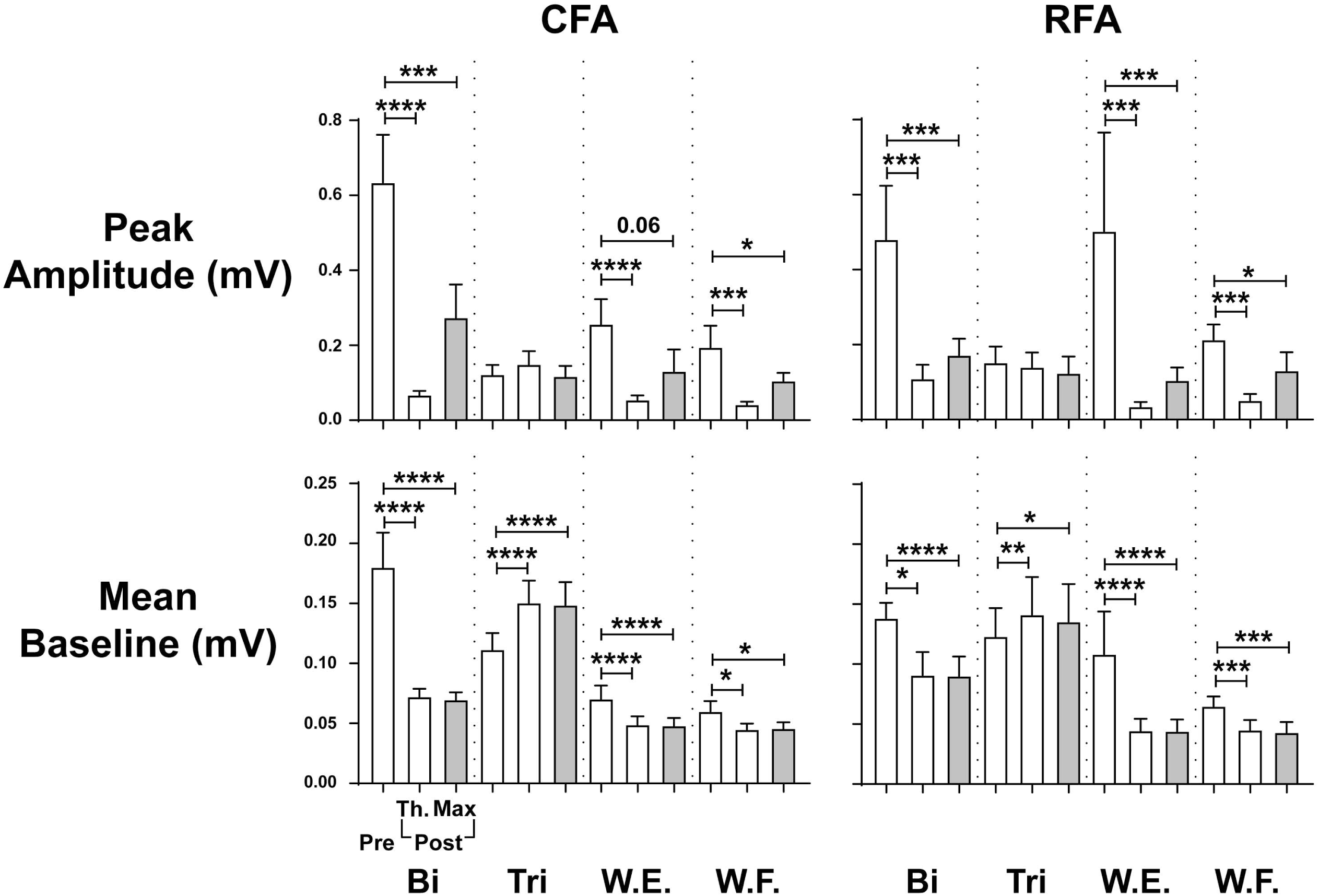
Abolished peak activity and baseline for forelimb muscles are not rescued when current threshold is increased. Bar graphs show reductions in StTA peak amplitude and mean baseline values as a function of MUS injection. Pre-peak amplitude measures in CFA and RFA obtained at threshold (white bars) are significantly reduced in biceps, wrist extensors and flexors (see **Table 1**). Using the highest applied current value for this experiment (80µA, gray bars), StTA were unable to revert to pre-levels. Similarly, pre-mean baseline measures in CFA and RFA are significantly reduced in all four muscles, and did not revert to pre StTA levels with 80µA currents.

MUS significantly reduced the percentage of activation from applied current pulses for all muscles in all rats in CFA (**Fig. 9**). On average, bicep activation was significantly reduced from 84% to 64% (F_(1,41.99)_ = 72.8644, p < 0.0001), wrist flexors were significantly reduced from 83% to 68% (F_(1,41)_ = 30.6232, p < 0.0001), wrist extensor activation was significantly reduced from 80% to 68% (F_(1,42.02)_ = 21.726, p < 0.0001), and tricep activation was significantly reduced from 72% to 61% (F_(1,42.15)_ = 37.005, p < 0.0001). Since StTA’s are an average of all stimulation trains that either induce or fail to induce muscle activation, these data suggest that much of the observed loss of peak amplitude can be attributed to average reductions in the likelihood of muscle activation. Moreover, the reductions in activation probability may, in part, explain the observed atypical ICMS-evoked movements post-MUS in CFA (see above section). When only selecting StTA values from sites where movement was evoked post-MUS (15/26 active sites), we still found a significant reduction in peak evoked amplitude in biceps, wrist flexors, wrist extensors (**Table 1**). Mean baseline amplitude for all muscles was also significantly reduced when looking at muscles from all sites and from only the active sites. These findings would suggest that the previous reductions in peak amplitude and mean baseline of StTAs were not simply a result of the non-active sites reducing the overall averages, but of a general reduction across all recorded CFA sites.

**Figure 9:**
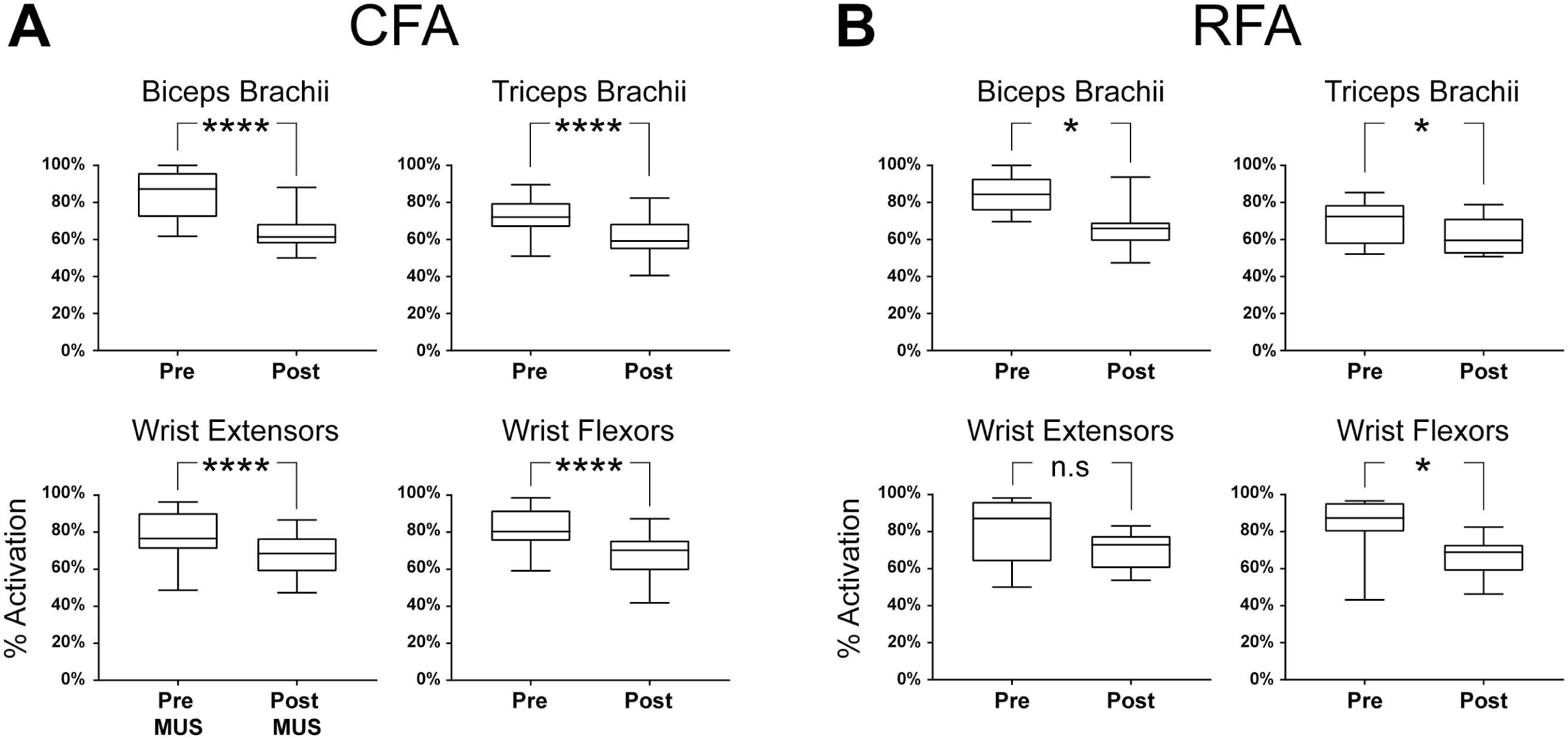
Some of the observed loss of forelimb muscle activity is due to lowered probability of muscle activation from CFA and RFA cortical stimulation. Box and whisker plots show the percentage of muscle activation during one-minute stimulation trials before and after MUS injection in CFA and RFA. Muscles were considered activated if individual EMG amplitude measures were 2.5 standard deviations from baseline. We found significantly reduced activation in each muscle when CFA was stimulated [biceps: F_(1,41.99)_ = 72.8644, p < 0.0001; triceps: F_(1,42.15)_ = 37.005, p < 0.0001; wrist extensor: F_(1,42.02)_ = 21.726, p < 0.0001; wrist flexors: F_(1,41)_ = 30.6232, p < 0.0001]. Muscle activation was also significantly reduced in biceps [F_(1, 8.839)_ = 9.0827, p < 0.05], triceps [F_(1,8.07)_ = 6.8884, p < 0.05], and wrist flexors [F_(1,8.96)_ = 8.65, p < 0.05] reduced when RFA is stimulated.

Like CFA, reductions in the StTA peak amplitude and mean baseline measures for RFA were observed following MUS inactivation (**Fig. 8**). The peak amplitude for StTAs were significantly reduced in biceps, wrist extensors and flexors (**Table 1**). Also, mean baseline amplitude values were significantly reduced for biceps, wrist extensors, wrist flexors, and triceps. On average, bicep activation was significantly reduced from 84% to 66% (F_(1, 8.839)_ = 9.0827, p < 0.05), wrist flexor activation was significantly reduced from 83% to 66% (F_(1,8.07)_ = 6.8884, p < 0.05), and triceps activation was significantly reduced from 67% to 62% (F_(1,8.96)_ = 8.65, p < 0.05; **Fig. 9**). Though there was an average reduction from 80% to 71%, wrist extensor activation was not found to significantly differ after MUS application. Interestingly, though earlier suppression of ICMS-evoked movement in RFA would suggest reduced muscle activation relative to CFA, and three out of the four muscle groups in RFA show a significant reduction in activation following MUS injection, stimulation to RFA was able to activate forelimb muscles at a similar probability to CFA (62% in RFA compared to 61% in CFA). Moreover, reductions in the StTA peak amplitude values are comparable between CFA and RFA – i.e. no significant differences in peak amplitude exists between biceps, triceps, wrist extensors, and wrist flexors, which suggest that loss of ICMS-evoked movements in RFA is not simply due to a loss of overall activation, but instead an inability to activate specific timing or patterning within CFA that is necessary to induce movement.

### Increases in applied current do not rescue muscle activity

Due to the data which show that intact threshold values may naturally increase as a consequence of MUS inactivation (see **Fig. 5**), we considered the possibility that the suppression of StTA amplitude and mean baseline for each muscle observed following MUS injection was due, in part or in total, to a general increase in the threshold necessary for activation, and that StTA values closer to pre-levels would be observed if applied ICMS currents were increased. To test, we compared the StTAs at threshold with the maximum applied ICMS current (80µA) after MUS injection (**Fig. 8**). We were still able to observe general and significant decrease in StTA peak amplitude in both CFA and RFA (**Table 1**), though the mean value was higher than those seen when currents were applied at threshold. In CFA, the peak amplitude for biceps and wrist flexors saw significant decreases when compared against pre-MUS amplitude, in a manner resembling post-MUS peak amplitude when currents were applied at threshold. Though not significantly different, wrist extensors also saw a general reduction in peak amplitude. Likewise, mean baseline values were still significantly reduced when applying maximum current for biceps, triceps, wrist extensors, and wrist flexors. RFA StTA peak amplitude measures were reduced in biceps, wrist extensors, and wrist flexors; Mean baseline was reduced in biceps, wrist extensors, and wrist flexors, triceps. Additionally, with the exception of wrist flexors which saw a moderate, but significant, average activation increase of 7.7% in CFA and 3.5% in RFA, the likelihood of activation did not significantly increase when compared against applied currents at threshold post-MUS, suggesting no meaningful improvement in either activation likelihood or intensity of muscle response to maximum applied current. Taken together, these results indicate that the observed loss of StTA activity is unlikely to be due to a general increase in threshold, but rather a direct consequence of MUS. Furthermore, the notion that the intact ICMS movements observed in CFA can be explained by an increase in required activation threshold is not sufficient enough to explain the how some CFA activity persists, but not in RFA.

## DISCUSSION

### Summary of Findings

Our work highlighted several features of the ipsilesional (pre)motor control of forelimb muscle when M1 is inactivated in rodents. We found that the inhibitory effects of MUS injections at CFA disproportionally impaired the indirect RFA sites when compared against direct CFA sites. Though both CFA and RFA activation of individual forelimb muscles were similarly impaired, ICMS-evoked movements are largely abolished in RFA while still present within CFA, and suggests that ICMS responses in RFA depend on intact CFA-RFA reciprocal connections (Rouiller et al., 1993) and resembles patterns of dependence found in the premotor-motor areas in non-human primates (NHP; Dum and Strick 2002, 2005; Schmidlin et al., 2008). In CFA sites where ICMS responses had persisted *post*-MUS, formally distal sites were likely to evoke proximal responses, suggesting that distal forelimb movements might be more sensitive to perturbations in CFA-RFA connectivity or to the increased GABAergic inhibition, relative to proximal sites. The presence of evoked movements during the *post*-MUS phase in only CFA but not RFA suggests that direct corticofugal projections from RFA to the brainstem and spinal cord are not sufficient to evoke movements without CFA, potentially hinting that the rodent forelimb motor pattern is largely encoded within CFA. Thus, the forelimb motor pattern is partially mediated by rodent motor cortex in a manner similar to serially processed premotor-motor interaction observed in the forelimb motor areas of NHPs, as also observed in recent rodent studies looking at forelimb control (Cerri et al., 2003; Shimazu et al., 2004; Schmidlin et al., 2008; Smith et al., 2010; Deffeyes et al., 2015; Kunori & Takashima, 2016).

Curiously, with the exception of the RFA in one rat (see face data point in Fig. 3), we observed that *post*-ICMS movements became largely suppressed in both RFA and CFA in a manner similar to MUS injections in NHP motor areas (Stepniewska et al., 2014). This contrasts with prior findings that showed *post* sites evoking non-forelimb (neck, face, vibrissae, hindlimb) movements at *pre*-like current thresholds in rodents (Okabe et al., 2016). As a result, while our face and trunk site area did not significantly increase following injection, our no response site area did. These no response sites appeared to neither correlate with distance from injection site nor prior threshold values yet do not appear to be stochastically distributed. Rather they appeared in higher quantities towards the caudal half of CFA (cCFA) and suggests histological differences in rostral (rCFA) vs cCFA. This result might differ from Okabe et al. (2016) might be a byproduct of experimental design, or a physiological difference between rodent strains (VandenBerg et al., 2008).

### A rationale for suppressed activation pattern within RFA and CFA

The observed pattern of GABAergic inactivation in CFA, where inactivation disproportionately affected cCFA hints at an underlying excitatory or inhibitory difference between the rCFA and cCFA. In NHPs, there exist a similar topography where PMv preferentially projects to rostro-lateral part of M1 (Dancause et al., 2006). Earlier work from our lab might suggest a similar connectivity via corticocortical between RFA and CFA, as: 1) rCFA injuries by controlled cortical impact trend towards greater RFA ICMS deficits compared to cCFA injuries (Nishibe et al., 2010); and, 2) we found some tract-tracing evidence that RFA to CFA projections are denser in rostral-most regions (Bury & Nudo, unpublished data). These findings could indicate a rostral bias for motor encoding, one that is only activated via direct artificial stimulation to rCFA but not indirect corticocortical fiber stimulation via RFA or cCFA. Given the similarity of EMG activity between the CFA and RFA post-MUS, an alternative and intriguing possibility could be that rodent forelimb movements are dependent on dual CFA and RFA output, whereby only the rCFA region contains sufficiently dense corticocortical connectivity to activate both CFA and RFA corticofugal fibers. Though this configuration would suggest an interaction between RFA and CFA dissimilar to NHP PM and M1, there has been some evidence that that these regions may differ, such as more diversity in RFA’s modulatory effects on CFA when compared against PM’s effects on M1 (Deffeyes et al., 2015, Cerri et al., 2003; Prabhu et al., 2009), or that more equivalent corticofugal projection patterns exist between rodent CFA and RFA than what is observed in NHP M1 and PM (Li et al., 1990; Dum and Strick, 1991; He et al., 1993; Rouiller et al., 1993; Borra et al., 2010; Saiki et al., 2014). These might hint at a heavier reliance on subcortical integration, and therefore greater dual RFA/CFA integration for forelimb movement in rodents.

### Differences between the organization of rodent and NHP motor representation

In NHPs it has been observed that specific ICMS-evoked movements are suppressed in premotor and supplementary motor areas when their corresponding M1 movements are inhibited, suggesting that these interconnected motor areas can be subdivided into functional domains (Stepniewska et al, 2014; Cooke et al., 2015). Here, the topographic layout of NHP forelimb motor maps revealed a feature of motor connectivity. In contrast, the mosaic pattern often observed in rodent forelimb motor maps lacks this topography. We still wondered whether proximal or distal sites in RFA connect with their respective sites in CFA. Or to put another way, do the loss of either proximal or distal movements have a higher likelihood of influencing their respective movement in RFA? Though the experimental parameters are fundamentally different, given that some distal sites evoked proximal movements post-MUS, intact CFA sites following injection had a higher likelihood of representing proximal movements, and RFA proximal movements were no less spared when compared to their distal counterparts, we would infer that movement representations in premotor area do not interconnect as specific functional domains like those observed in NHPs, and instead might depend on the complete integrity of the entire motor area, in sharp contrast to NHPs.

### Comparisons of Observed EMG Response to other work

The data presented here contributes to the limited number of papers looking at forelimb EMG by cortical activation in rats (Liang et al., 1993; Brus-ramer et. al, 2009; Deffeyes et al., 2015). Though our average EMG latency values obtained for all muscles in all rats was (∼15-17ms) differed by the ∼9.6ms observed in Liang et al. (1993), we found a similar distribution of latency values to those of Deffeyes et al. (2015). Both CFA and RFA values had similar latencies for all four muscle groups using the parameters of our experiment. Unlike prior studies, we did not exceed stimulation values of 80µA but were able to elicit similar EMG latency values with 3-pulse stimulation, indicating that temporal features of rodent EMG responses are conserved in the presence of differing stimulation parameters. Contrasted with NHP PM, which are unable to evoke EMG responses in the absence of paired-M1 activation (Cerri et al., 2003), we found that we were able to evoke EMG responses with solely RFA at a maximum amplitude that was not significantly different from CFA and in a manner similar to the single pulse stimulation observed in Liang et al. (1993), further hinting at potential equivalent corticospinal connectivity from RFA and CFA to spinal cord in rats compared to NHP, though to our knowledge no studies that have delineated any difference between rodent and primate corticofugal connections if one such exists.

## Conclusions

The results here are a first step in understanding how injury without damage to cortical tissue effects recovery, in contrast to traditional forms of injury (stroke and TBI) where inactivation is elicited by cortical damage. It was important to understand the acute consequence of temporary ‘injury’ to motor cortex and subsequent forelimb activity before attempting to disentangle adaptive/maladaptive responses during the (sub)acute stages of injury when compared to traditional stroke or TBI models. Our finding that RFA appears functionally ineffective, but not ‘inactive’ when considering EMG data comparisons between CFA and RFA could hint at several interesting possibilities. Reorganization (subacute or otherwise), that reinstates RFA function could be a natural consequence of a permanent injury model, while inactivation (with the preservation of brain tissue) may prove to be epiphenomenal or even maladaptive. Follow-up inactivation studies need to be done to directly look at the ipsilesional MUS effects over longer time-scales to see if these effects are consistent with those observed in more permanent injuries. One study showed that at the very least, the functional loss of activity – but not tissue – in the contralesional hemisphere produces observable changes in the recovery of the ipsilesional hemisphere, suggesting that adaptive reorganizational processes that result from loss of a brain area do not require tissue loss, but loss of activity, and might extend to ipsilesional reorganizational processes (Mansoori et al., 2014).

